# A member of the ferlin calcium sensor family is essential for *Toxoplasma gondii* rhoptry secretion

**DOI:** 10.1101/304048

**Authors:** Bradley I. Coleman, Sudeshna Saha, Seiko Sato, Klemens Engelberg, David J. P. Ferguson, Isabelle Coppens, Melissa B. Lodoen, Marc-Jan Gubbels

## Abstract

Invasion of host cells by apicomplexan parasites such as *Toxoplasma gondii* is critical for their infectivity and pathogenesis. In *Toxoplasma*, secretion of essential egress, motility and invasion-related proteins from microneme organelles is regulated by oscillations of intracellular Ca^2+^. Later stages of invasion are considered Ca^2+^-independent, including the secretion of proteins required for host cell entry and remodeling from the parasite’s rhoptries. We identified a family of three *Toxoplasma* proteins with homology to the ferlin family of double C2 domain-containing Ca^2+^ sensors. In humans and model organisms such Ca^2+^ sensors orchestrate Ca^2+^-dependent exocytic membrane fusion with the plasma membrane. One ferlin that is conserved across the Apicomplexa, TgFER2, localizes to the parasite’s cortical membrane skeleton, apical end, and rhoptries. Unexpectedly, conditionally TgFER2-depleted parasites secreted their micronemes normally and were completely motile. However, these parasites were unable to invade host cells and were therefore not viable. Specifically, knockdown of TgFER2 prevented rhoptry secretion and these parasites failed to form the moving junction on the parasite-host interface necessary for host cell invasion. Collectively, these data demonstrate that the putative Ca^2+^ sensor TgFER2 is required for the secretion of rhoptries. These findings provide the first regulatory and mechanistic insights into this critical yet poorly understood aspect of apicomplexan host cell invasion.

**Graphical abstract:** 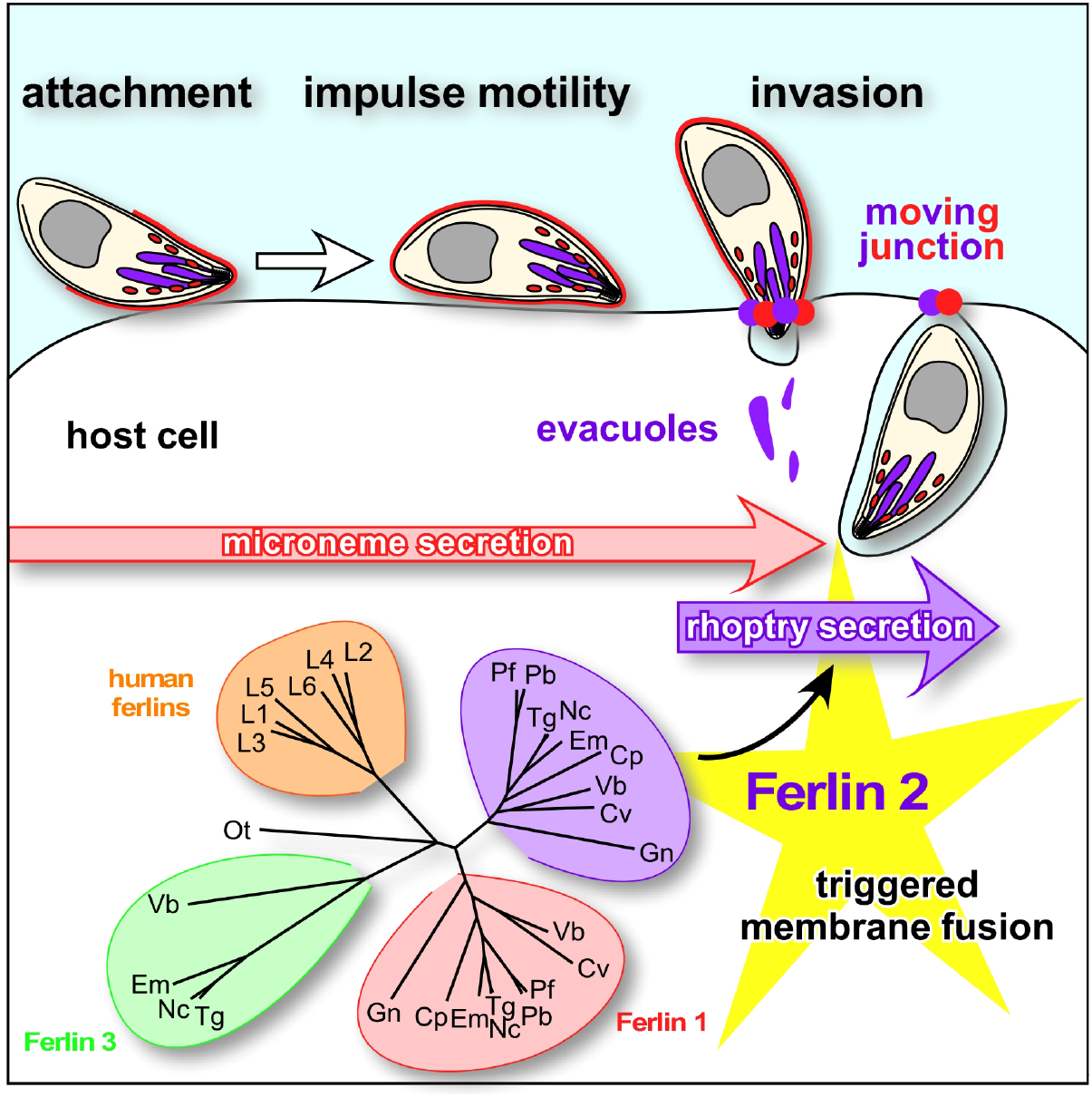

## Introduction

The apicomplexan parasite *Toxoplasma gondii* infects one in every three humans. Clinical symptoms of toxoplasmosis derive from the tissue destruction and inflammation caused by repeated rounds of host cell invasion, intracellular replication, and lytic egress of the tachyzoite life stage. Egress is mediated by intracellular Ca^2+^ [Ca^2+^]_i_ fluctuations that trigger release of proteins from the microneme organelles (1). Following egress, parasites move via gliding motility to a new host cell, triggered by additional parasite [Ca^2+^]_i_ oscillations facilitating further release of micronemes (2). Subsequent host cell invasion also relies on micronemal proteins (3). The micronemes are localized at the apical end of the parasite, and are released from the apical tip (4). Following initial recognition of a host cell, the parasite engages in a tighter interaction with the target cell, mediated by proteins secreted from the rhoptries. The club-shaped rhoptries are anchored at parasite’s apical end from where they secrete their contents (5, 6). Proteins in the apical rhoptry neck (RONs) are secreted into the host cell before the proteins residing in the more basal rhoptry bulb (ROPs) are released (7). RONs function in tightening the parasite:host attachment interface by forming a moving junction (MJ) whereas ROPs modulate a variety of host cell pathways to accommodate intracellular replication (6). Rhoptry secretion must be preceded by microneme secretion and requires recognition of a host cell, but their release is generally assumed to be Ca^2+^-independent although the molecular details of the underlying signal transduction pathways and the mechanism of rhoptry exocytosis remain obscure (10). Both the micronemes and rhoptries are conserved across Apicomplexa and are key to the intracellular life style of these pathogens (9). The last step in establishing infection of a new cell is secretion of host cell-remodeling proteins from the parasite’s dense granules, which again is believed to be a Ca^2+^-independent process.

The Ca^2+^ signal during egress and invasion is transduced by several molecular mechanisms, including calmodulin (2), calcineurin (11), Ca^2+^-dependent protein kinases (CDPK1 (12), CDPK3 (13–15)), and at the point of microneme exocytic membrane fusion by the DOC2.1 protein (16), referred to as TgDOC2 hereafter. The organization of TgDOC2 is unusual compared to well-studied Ca^2+^-triggered exocytosis models since the only identifiable domain in this large protein is the namesake double C2 domain (“DOC2”). In model organisms coordination between at least three DOC2 domain proteins execute the Ca^2+^-mediated vesicle fusion with the plasma membrane typically in combination with a transmembrane domain in one or two proteins (18, 21–23). Ca^2+^ exerts its function through association with positionally conserved Asp residues in C2 domains, which then associate with membrane or other proteins facilitating membrane fusion (17, 18). The ferlins make up a unique branch of the DOC2 domain protein family tree as they contain five to seven C2 domains rather than two, and they are relatively large (200–240 kDa). The ferlins comprise an ancient eukaryotic protein family present in most protozoa (except amoeba and fungi) including the Apicomplexa and all metazoa (except higher plants) (19). Although ferlins are not expressed in neurons and lacking in yeast they are relatively understudied, they typically function in membrane fusion, vesicle trafficking and membrane repair, Dysfunction of human ferlins can cause deafness and muscular dystrophy (20).

To better understand the machinery underlying *Toxoplasma* Ca^2+^-mediated exocytosis we evaluated the DOC2 domain family in the Apicomplexa. Next to TgDOC2.1 (16) we identified three members of the ferlin family, two of which are widely conserved across Apicomplexa. We determined that TgFER2, the most-conserved representative, is essential for host cell invasion and required for rhoptry secretion. These findings provide critical insight into the poorly understood mechanisms of rhoptry secretion while raising the possibility that, contrary to common assumptions, rhoptry secretion is a Ca^2+^-dependent process.

## Results

### The *T. gondii* genome encodes three ferlin proteins

Next to TgDOC2, A series of BLAST searches of the *Toxoplasma* genome identified four additional proteins containing two or more C2 domains of which three also contained a transmembrane domain. Two proteins had clear homology to the ferlin family of Ca^2+^ sensitive membrane fusion proteins (Fig. 1A). We named these proteins TgFER1 (TGME49_309420) and TgFER2 (TGME49_260470). The other DOC2 proteins, TGME49_295472 and TGME49_295468, are adjacent in the genome but these are annotated as a single gene in the ontological region in *Neospora* and *Eimeria spp*. This merged protein also possesses the global architecture of a ferlin, and was named TgFER3, but it diverges from the family by its extensive degeneration of C2 domains and unusual length: with 2670 amino acids TgFER3 is 40% longer than the other ferlins (Fig. 1A).

**FIG 1.**
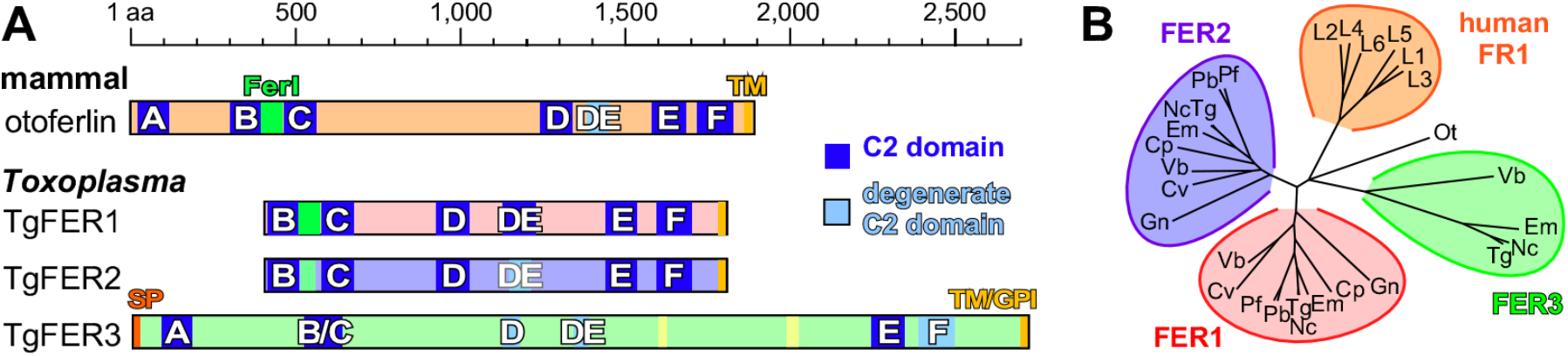
**(A)** *Toxoplasma* encodes three ferlin proteins and human otoferlin is shown for comparison. Ferlins are defined by 5 to 7 C2 domains (labeled A-F) and a C-terminal transmembrane (TM) domain and typically a ‘FerI’ domain of unknown function. TgFER3 contains an N-terminal signal peptide, which, in combination with the C-terminal TM domain, could signal GPI-anchor addition at the C-terminus. Yellow shades in TgFER3 represent coiled-coil domains. (**B**) Phylogenetic analysis of apicomplexan, chromerid and human ferlins. The following abbreviations are used: human ferlins “L1-L5” FR1L1-5 (FR1L1 (dysferlin; O75923.1), FR1L2 (otoferlin; Q9HC10.3), FR1L3 (myoferlin; Q9NZM1.1), FR1L4 (A9Z1Z3.1), FR1L5 (A0AVI2.2), FR1L6 (Q2WGJ9.2)), Ot: green algae *Ostreococcus tauri* (Q01FJ7); Chromerids “Vb” *Vitrella brassicaformis* (VbFER1 (Vbre_12074 + Vbra_12075), VbFER2 (Vbra_9198)) and “Cv” *Chromera velia* (CvFER1 (Cvel_17519.2) and CvFER2 (Cvel_9223)); Apicomplexa “Tg” *Toxoplasma gondii* (TgFER1 (TGME49_309420), TgFER2 (TGME49_260470), TgFER3 (TGME49_295472 + TGME49_295468)) “Nc”, *Neospora caninum* (NcFER1 (NCLIV_053770), NcFER2 (NCLIV_026570), NcFER3 (NCLIV_002280)), “Em” *Eimeria maxima* (EmFER1 (EMWEY_00002120), EmFER2 (EMWEY_00009280), EmFER3 (EMWEY_00017650)), “Pf” *Plasmodium falciparum* (PfFER1 (PF3D7_0806300), PfFER2 (PF3D7_1455600)), “*?VTlasmodium berghei* (PbFER1 (PBANKA_122440), PbFER2 (PBANKA_131930)), “Cp” *Cryptosporidium parvum* (CpFER1 (cgd8_2910), CpFER2 (cgd2_2320)), “Gn” *Gregarina niphandrodes* (GnFER1 (GNI_063830), and GnFER2 (GNI_073830)). Alignment and unrooted Jules-Cantor phylogenetic tree were generated in Geneious (v.6.1.6) (63)) from a MUSCLE alignment using neighbor-joining. Note that the FER1 and FER2 nodes for Tg and Nc are barely discernable at this scale.

Using human otoferlin as a reference, the C2 domains in *T. gondii* ferlins 1–3 follow the typical paired C2 pattern (Fig. 1A). The absence of the C2A domain in TgFER1 and TgFER2 is not unusual as this domain is missing in the majority of studied ferlins (19). All ferlins studied to date contain the FerI domain of as yet unknown function, which is present in TgFER1, slightly degenerate in TgFER2, and undetectable in TgFER3. We queried the conservation of ferlins in representative apicomplexan organisms and their closest free-living relatives, the Chromerids (24). Clear orthologs of TgFER1 and TgFER2 were universally present, but TgFER3 orthologs were restricted to the Coccidia *(Neospora, Sarcocystis, Eimeria)* and somewhat surprisingly, to the chromerid *Vitrella brassicaformis* (Fig. 1B). This suggests that TgFER3 was present in the last common ancestor of Chromerids and Apicomplexa but was lost from all apicomplexan lineages except the Coccidia.

### TgFER2 is essential for completing the lytic cycle

The most widely studied Ca^2+^-mediated process in *Toxoplasma* is the release of microneme proteins. Given the documented roles of DOC2 and ferlins in Ca^2+^-mediated secretion we hypothesized that apicomplexan ferlins are involved in microneme secretion. To test this hypothesis we probed the function of TgFER2 by replacing its promoter with a tetracycline regulatable promoter (25) and simultaneously inserted a single N-terminal Myc epitope to provide localization data (Fig. 2A). The genotype was validated by diagnostic PCR (Fig. 2B). Western blots of total parasite lysates probed with α-Myc antibodies marked a single protein consistent with the 160 kDa predicted molecular weight of TgFER2 (Fig. 2C). Regulation of TgFER2 was demonstrated by exposing FER2-cKD parasites to anhydrous tetracycline (ATc) for 48 hours to block TgFER2 transcription. Myc-TgFER2 was undetectable by western blot (Fig. 2C) or IFA (Fig. 2D) confirming efficient protein knock-down. TgFER2-depleted parasites did not form plaques after 7 or 14 days (Fig. 2E). No observable changes in the morphology or growth rate of intracellularly replicating parasites was observed (see Fig. S1 in the supplemental material). TgFER2 therefore does not function in cell division or replication but is essential for completion of *Toxoplasma’*s lytic cycle.

**FIG 2.**
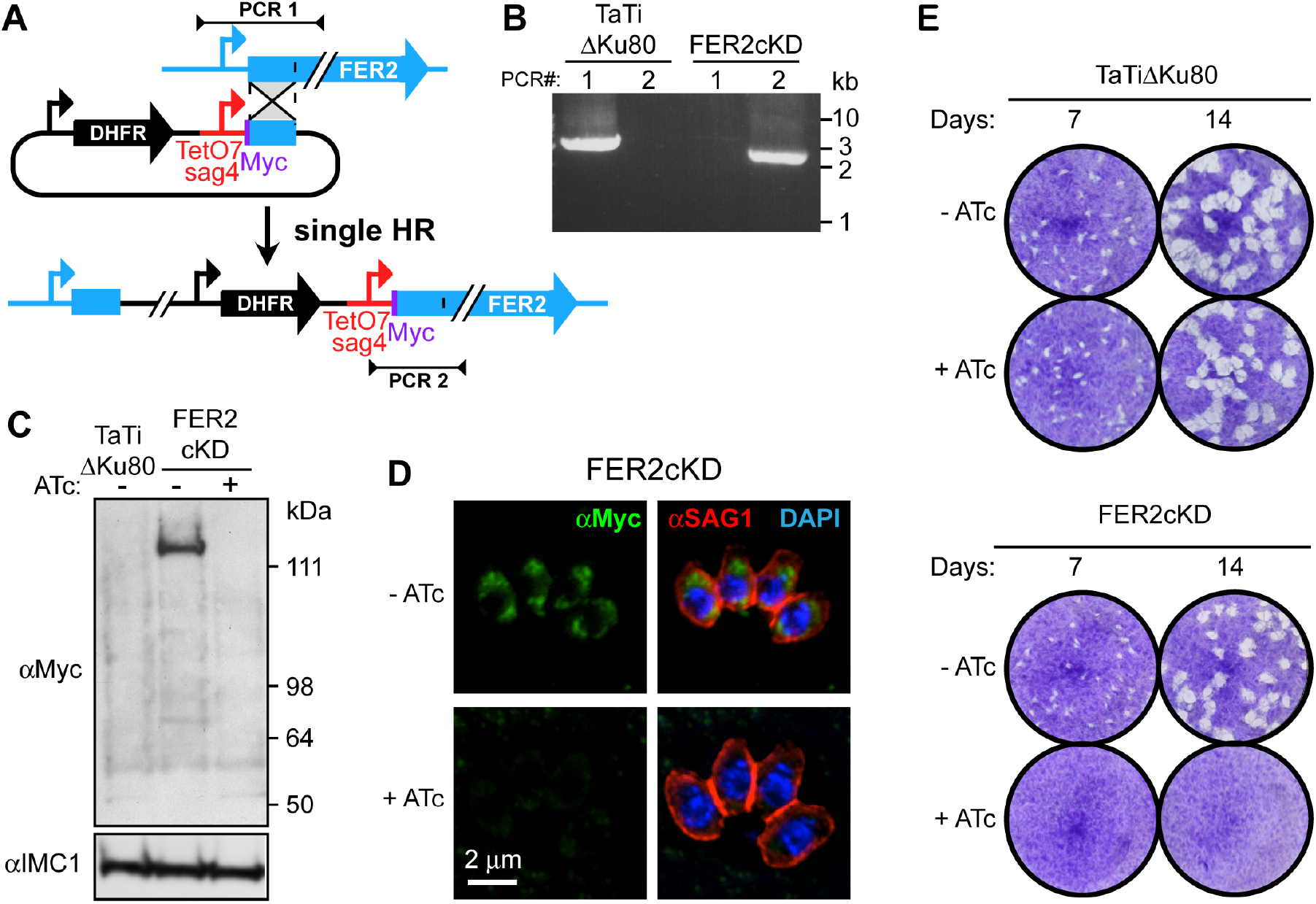
Generation and validation of a TgFER2 conditional knock-down parasite. (A) Schematic representation of single homologous promoter replacement with the anhydrous tetracycline (ATc) regulatable promoter TetO7sag4. Note that a Myc-epitope tag is simultaneously added on the N-terminus. Sites of diagnostic primer pairs used in panel B are indicated. **(B)** Diagnostic PCR of the parent line (TaTiΔKu80) and the FER2-ckD promoter replacement line using the primer pairs depicted in panel A. **(C)** Western blot demonstrating the conditional expression of the Myc-tagged TgFER2 allele. TgFER2 is downregulated to undetectable levels after 48 hr of ATc treatment. α-IMC1 is used as a loading control. **(D)** Immunofluorescence demonstrating the loss of Myc-TgFER2 expression upon ATc treatment for 20 hr. Parasites were fixed with 100% methanol. DAPI labels DNA and α-SAG1 marks the plasma membrane. (e) Plaque assays of parent (TaTiΔKu80) and FER2-ckD lines ±ATc treatment for times as indicated. No plaques are observed upon loss of TgFER2 expression.

### TgFER2 localizes on the IMC, rhoptries and inside the conoid

Myc-TgFER2 localization by IFA revealed a dispersed pattern not reminiscent of any defined *Toxoplasma* feature (Fig. 2D). Since the transmembrane domain is predictive of membrane association, we resolved the localization by immunoelectron microscopy (IEM). In intracellular parasites a comparatively small number of gold particles were distributed throughout the cytoplasm (Fig. 3A). Gold was notably enriched at the cytoplasmic side of the inner membrane complex (IMC) (Figs. 3A, B) and within the internal structures of the conoid at the apical end (Figs. 3A, C-E). Since intracellular parasites do not actively secrete micronemes, the TgFER2 localization might be different in extracellular parasites that are actively secreting micronemes. In extracellular parasites we indeed observed a different pattern with gold particles patched on the cytoplasmic side of the rhoptries (Fig. 3F-H). Critically, TgFER2 labeling was not specifically observed on the micronemes in either intracellular or extracellular parasites, which appeared to be inconsistent with our initial hypothesis of a role for FER2 in microneme secretion.

**FIG 3.**
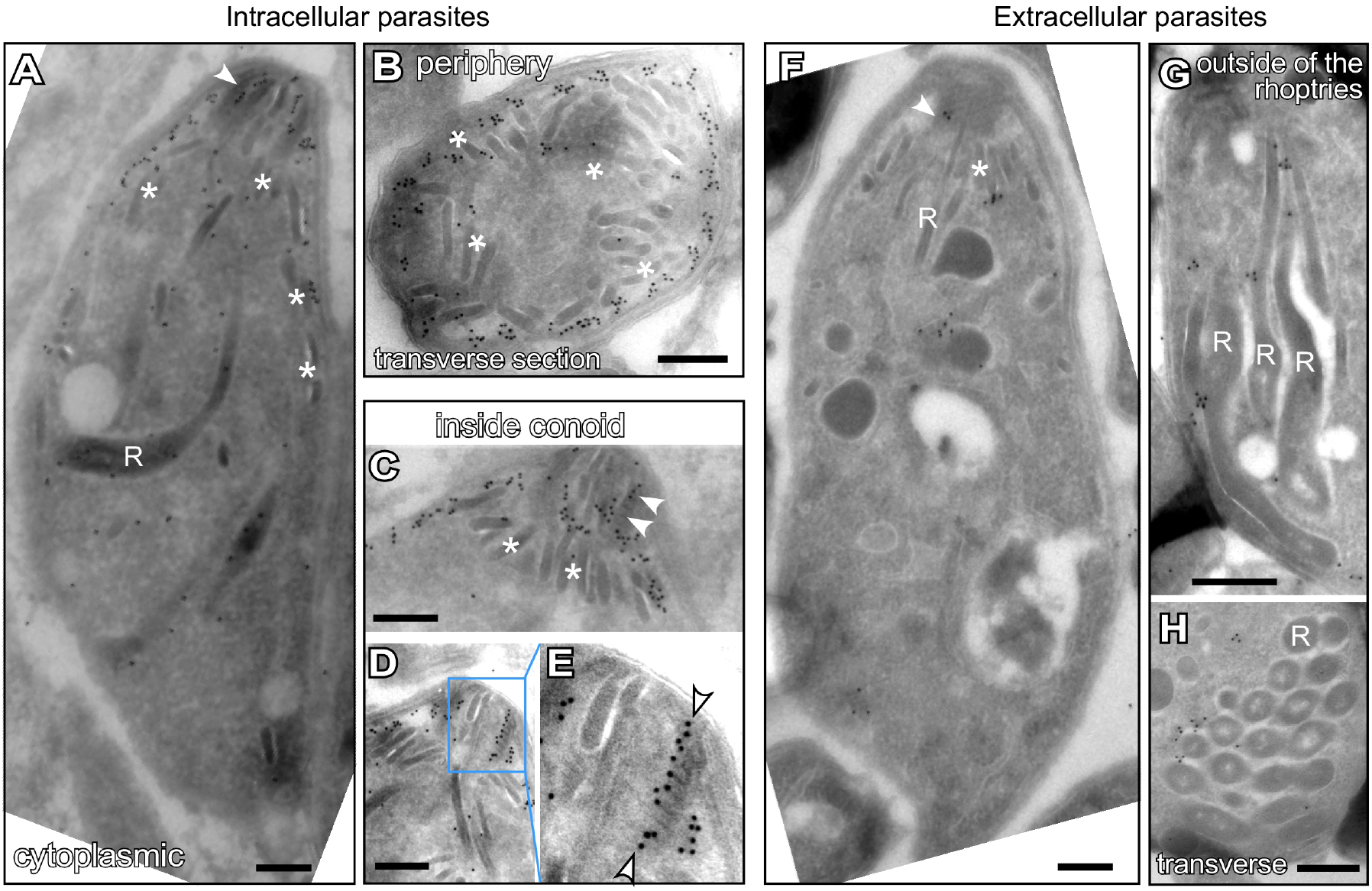
Subcellular localization of TgFER2. Immunoelectron microscopy of intracellular (**A-D**) and extracellular (**F-H**) *Toxoplasma* tachyzoites expressing an N-terminal Myc-epitope tagged TgFER2 from the endogenous locus under the TetO7sag4 promoter. In Intracellular parasites Myc antibodies direct gold particle clusters to the cytoplasmic side of the IMC and a strong enrichment inside the conoid, but no strong association with either micronemes and minor association with the rhoptries. In extracellular parasites gold particles are predominantly observed in clusters on the cytoplasmic side of the rhoptry membranes next to localization inside the conoid. R marks the rhoptries; asterisks mark micronemes, arrowheads mark gold beads in the conoid. Panel E is a magnification of the region marked in panel D. Scale bars are 250 nm.

### TgFER2 is not required for microneme secretion and conoid extrusion

To determine the lethality of TgFER2-depleted parasites we first assayed parasite egress. After 48 or 96 hours of ATc treatment, FER2-ckD parasites egressed normally when treated with Ca^2+^ ionophore A23187 (see Fig. S2 in the supplemental material). This suggests that the micronemes are secreted normally. The morphology and distribution of micronemes in TgFER2-depleted parasites is also normal by IFA and TEM (see Fig. S3 in the supplemental material). Since TgFER2 is present in the conoid we also examined conoid extrusion as another Ca^2+^-regulated process (26). Fig. S4 in the supplemental material shows that conoid extrusion is normal in the TgFER2 mutant.

Next we directly tested microneme protein secretion through Mic2 release (27). Both untriggered, low-level constitutive secretion and Ca^2+^ ionophore-induced microneme secretion occurred normally in the absence of TgFER2 (Fig. 4A, B). It is now apparent that micronemes are not uniform and that distinct populations containing distinct proteins exist within the parasite (28). We therefore reasoned that TgFER2 might act differentially on these populations and that this might explain our observations. Mic2 is secreted from a Rab5a/c-dependent population of micronemes. Another component of this population, Mic10, was also secreted normally in TgFER2-depleted parasites (Fig. 4A). To examine secretion from the Rab5a/c-independent population containing Mic proteins 3, 5, 8 and 11 we first assayed Mic8 secretion by western blot. This also proceeded normally in the absence of TgFER2. Labeling of Mic3, 5 and 8 by IFA on non-permeabilized parasites (29, 30) also confirmed secretion of these proteins to the surface of both FER2-replete and -depleted parasites (Fig. 4C, see Fig. S5A, B in the supplemental material). Thus, secretion of all micronemes is TgFER2 independent.

**FIG 4.**
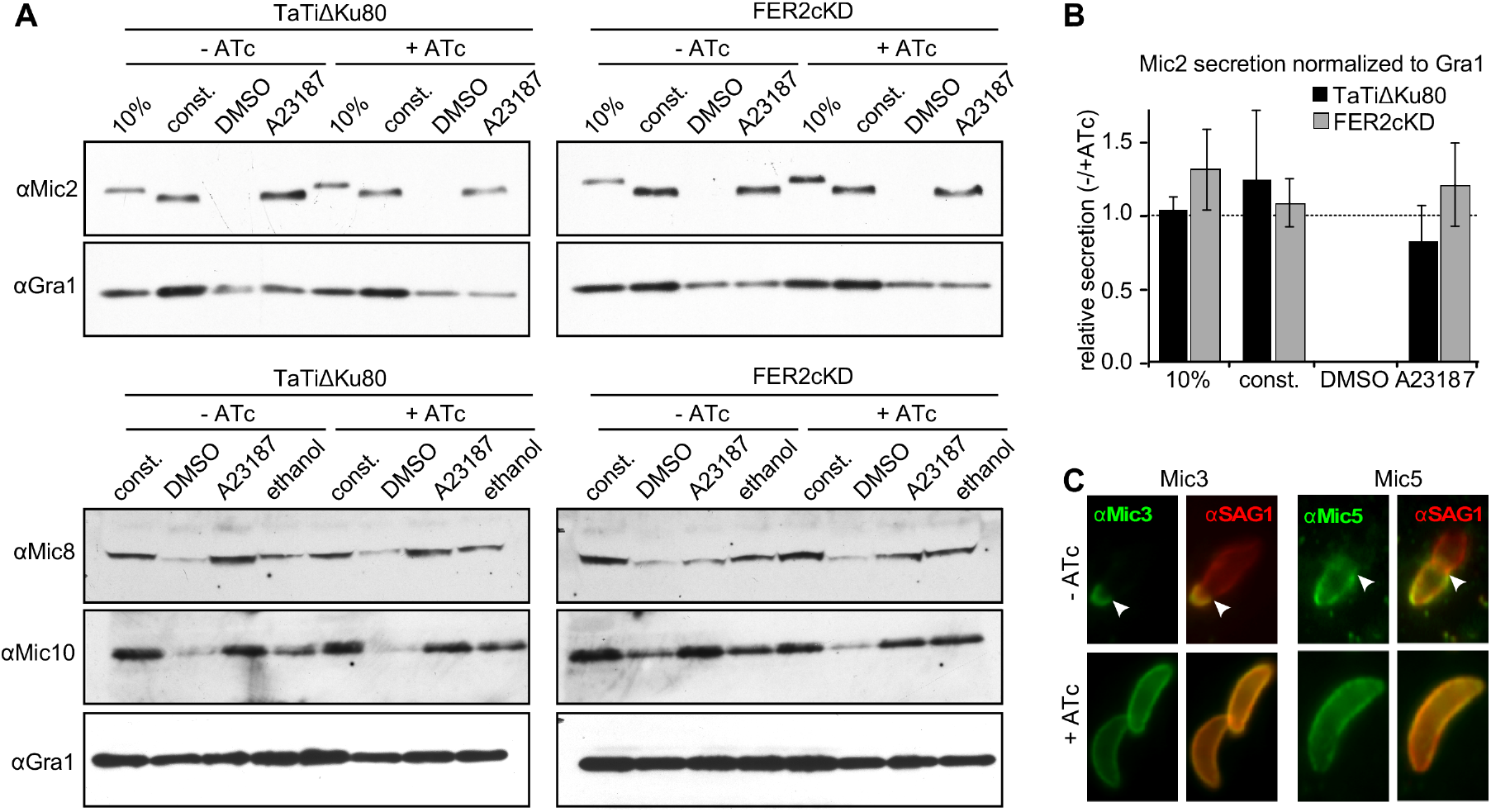
Microneme secretion of TgFER2 depleted parasites. (A) Microneme secretion assay by western blot. The lane labeled “10%” shows total parasite lysate corresponding with 10% of the parasites used in the secretion assay; const. represents constitutive secretion of extracellular parasites for 1 hr; A23187 and DMSO represent induced secretion with Ca^2+^ ionophore (1 μM A23187) and the vehicle control for 5 min. Ethanol represents 1% ethanol as trigger for microneme secretion. Microneme secretion of the classic population is detected by western blotting with α-Mic2, which shows a size shift upon secretion, and with α-Mic10. Secretion of the Mic3/5/8/11 microneme population is monitored with α-Mic8. α-Gra1, which detects dense granule secretion, is used as control. (**B**) Quantitation of Mic2 secretion normalized to GRA1 secretion shown in panel A. n=3 ±stdev. No statistical differences detected. (**C**) Secretion of the Mic3/5/8/11 population monitored by IFA using α-Mic3 and α-Mic5 (α-Mic8 data in Fig. S5A, B in the supplementary material). Extracellular FER2-ckD parasites ±ATc were placed on HFF cells. Host cells were permeabilized by 0.02% saponin (parasites are not permeabilized in this condition) so that only secreted Mic is detected. α-SAG1 marks the plasma membrane. Arrowhead marks the site of invading parasites at the boundary where the apical end of parasites is already inside the host cell and stripped of nearly all Mic and most SAG1 protein. Single color and phase panels are shown in Supplementary Fig. S5.

Surface antigen SAG1 and Mic8 were deposited in trails behind parasites ±ATc, implying that TgFER2-depleted parasites are still motile (see Fig. S5A, B in the supplemental material). This was confirmed by scoring the total number of motile parasites and the type of motility displayed by individual FER2 knockdown parasites by video microscopy (see Fig. S5C in the supplemental material). However, invading parasites, which are clearly identifiable by the stripping of Mic proteins off the apical invading parasite surface, were only observed in the presence of TgFER2 (Fig. 4C, see Fig. S5A in the supplemental material), suggesting an invasion defect independent of the micronemes.

### TgFER2 is required for host cell invasion

We further examined host cell invasion of TgFER2-cKD mutants through a series of invasion and attachment assays. As controls, we used mutants with defects at different points of host cell attachment and/or invasion. These include the TgDOC2 temperature sensitive (*ts*-DOC2) mutant devoid of all microneme secretion (16), the calcineurin (CnA) mutant, which secretes micronemes but does not attach properly (11), the AMA1 mutant, which secretes micronemes but shows an increase in aborted invasions due to failures in functional MJ formation (31, 32), and the DHHC7 mutant, which lacks the palmitoyltransferase responsible for anchoring the rhoptries at the apical end of the parasites and as a result is defective in rhoptry secretion (5).

Early events in parasite attachment are mediated by the binding of SAG proteins to glycans on the host cell surface. We assayed this by attachment of parasites to fixed host cells (29). Only the *ts*-DOC2 mutant demonstrated reduced attachment relative to the wild-type and uninduced controls, which indicates normal microneme secretion for all other mutants (Fig. 5A). Next we tested attachment and invasion by the “red-green invasion assay” (33). TgFER2 depleted parasites invaded at a much lower frequency (Fig. 5B). The defect intensified >3-fold upon prolonged TgFER2 depletion (96 hrs), which importantly did not affect their viability as these parasites are still fully capable of egress (see Fig. S2 in the supplemental material). As expected, all other mutants showed severe invasion defects. In this assay, quantifying the total number of parasites per field allows for an estimation of parasite attachment. By this metric, a defect in the attachment of TgFER2-depleted parasites to host cells was observed. By 96 hr of knockdown, the numbers of parasites attached to host cells dropped nearly 4-fold (Fig. 5B). This approaches the levels observed in the *ts*-DOC2 mutant where attachment and invasion are both severely defective. The similarity between the *ts*-DOC2 mutant, where the primary defect is in attachment, and the DHHC7 mutant, with a primary invasion defect, highlights that this assay is unable to distinguish between these two interconnected phenotypes.

**FIG 5.**
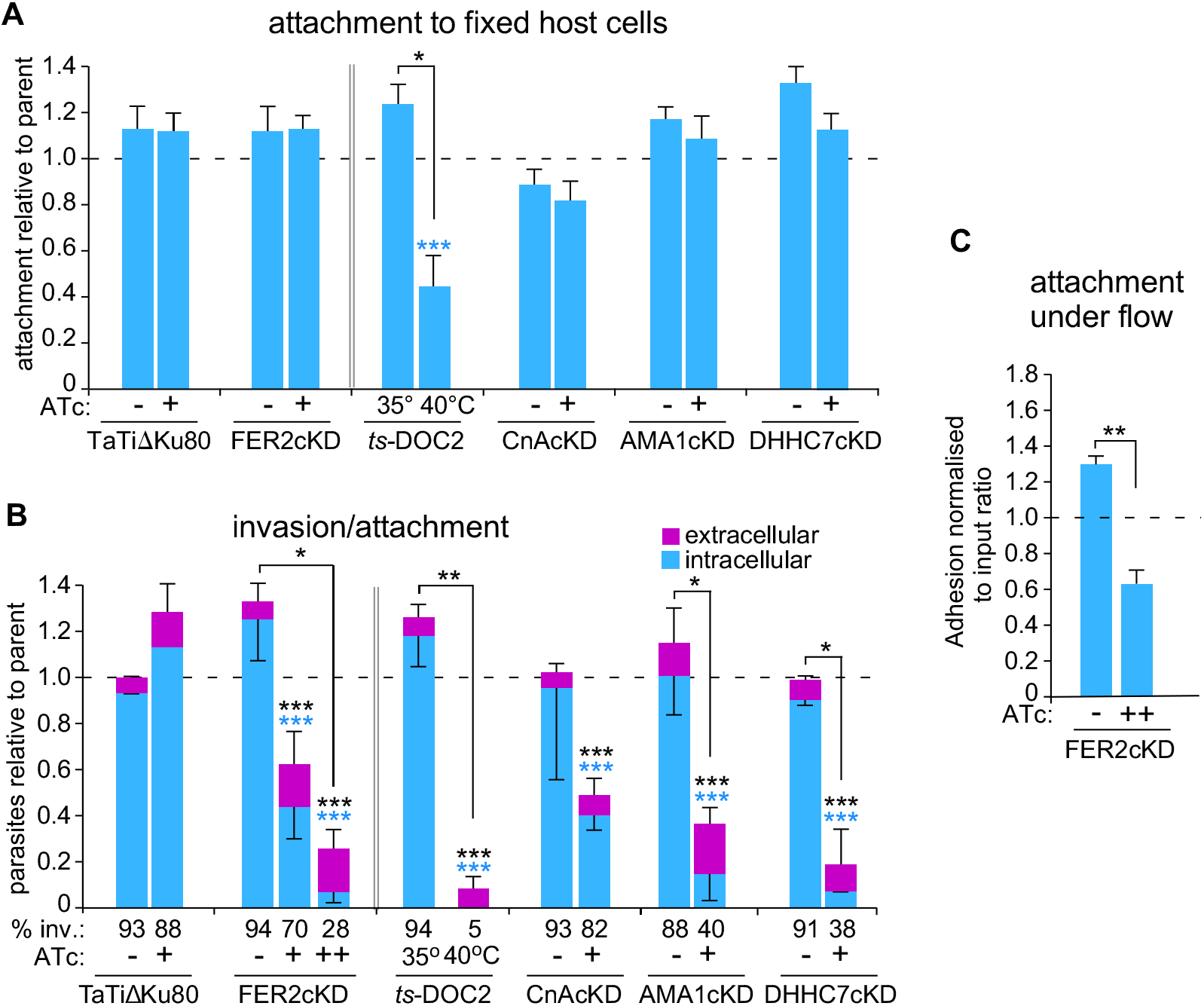
Invasion and attachment of TgFER2 depleted parasites. (A) Attachment to fixed HFF cells. Parasites as indicated were mixed 1:1 with the internal control (TaTiΔKu80 parasites expressing cytoplasmic YFP) and exposed to fixed HFF cells. All parasites were stained with α-SAG1 and control vs. test parasites counted; data are expressed relative to the internal control of TaTiΔKu80 -ATc (for *ts*-DOC2 TaTiΔKu80 was used for comparison rather than its direct parent). Mutants were induced with ATc for 48 hr except *ts*-DOC2, which was induced at 40°C for 48 hr. The dotted line represents the internal control level. n=5 +sem. Across samples statistics – or + ATc by one way ANOVA *** P<0.0001; Pairwise ±ATc statistics: Student’s *t*-test * P=0.014. (B) ‘Red-green invasion assay’ to determine invasion and attachment efficiency. Extracellular parasites were differentially stained from intracellular parasites with Alexa488 conjugated α-SAG1 before fixation; all parasites were subsequently stained following fixation and permeabilization with Alexa594 conjugated α-SAG1. For FER2, presence of ATc marked with “+” reflects 48 hr; “++” reflects 96 hr. n=3 + or – stdev. Across samples statistics – or + ATc: one way ANOVA; colored asterisks represent the variable compared across samples; black asterisks represent the total number of parasites. The % of invaded parasites is indicated at the bottom. Pairwise ±ATc statistics: Student’s *t*-test. * P<0.01, ** P=0.001, *** P<0.0001. (c) Parasite attachment to HUVECs under fluidic shear stress. Adhesion of the FER2cKD line ±ATc (96 hr) was compared. Parasite adhesion normalized to the ratio of each parasite population introduced into the fluidic channel is shown, wherein a value of 1.0 represents equivalent adhesion of the two populations. n=3 + stdev. **P<0.01 (Student’s t-test). See Supplementary Fig. S6 for additional controls.

It has been observed that the motility and attachment dynamics of *Toxoplasma* are different under conditions of shear stress in a flow chamber (34). To investigate whether these conditions might better clarify the phenotype of the TgFER2 knockdown, we measured the ability of FER2-cKD parasites ±ATc to adhere to human vascular endothelial cells (HUVEC) under flow. Depletion of TgFER2 led to a significant decrease in the number of parasites retained in the chamber (Fig. 5C and see Fig. S6 in the supplemental material), though attachment of TgFER2-depleted parasites was less compromised under flow relative to static conditions. This again demonstrates that TgFER2 is essential for invasion but does not pinpoint the nature of the defect. FER2-cKD parasites were then scrutinized for their interactions with host cells by video microscopy. Both wild type and FER2-ckD parasites were able to glide across the host cells. In contrast to control parasites, TgFER2 depleted parasites were not able to invade host cells (Fig. 6 and see Movie S1 in the supplemental material). Surprisingly, FER2-ckD parasites still exhibited “impulse motility” characteristic of invading parasites (35). This typical burst of forward motion immediately preceding invasion is followed by a momentary pause when parasites secrete the RONs and create the MJ before proceeding with invasion and parasitophorous vacuole formation. Both control and induced FER2-cKD parasites displayed bursts of impulse motility followed by a pause. However, only in control -ATc parasites this pause was followed with forward motion (invasion) at approximately half the original speed. In contrast, the velocity of +ATc parasites dropped essentially to 0 μm/sec and failed to invade. This observation suggests that TgFER2 functions in the very late stages of invasion and indicates that either the MJ is not formed, or is not of sufficient strength to support the force required for parasite penetration into cells.

**FIG 6.**
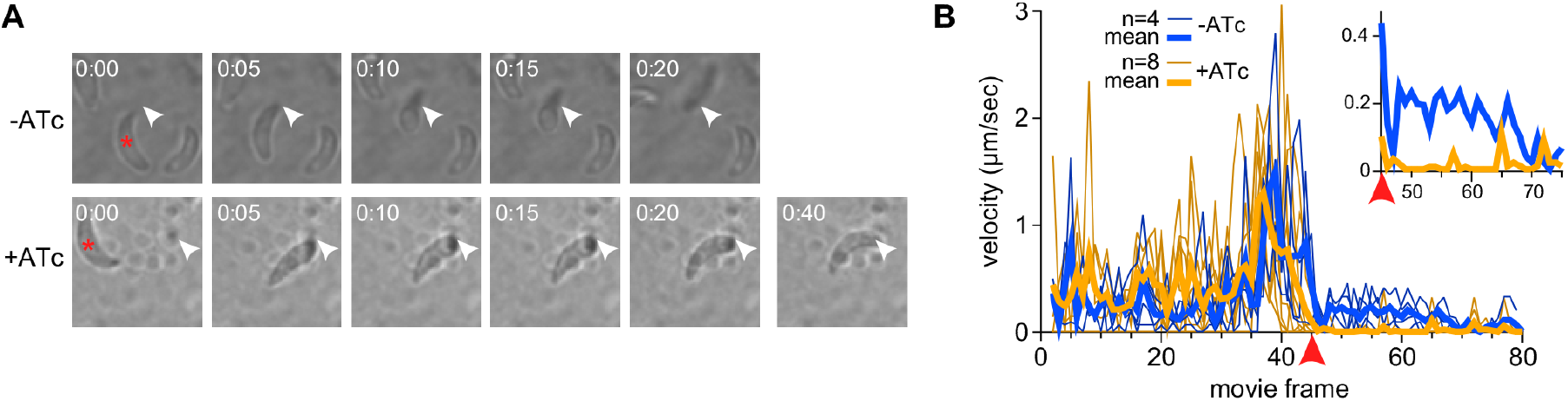
Impulse motility and host cell invasion. **(A)** Still panels from movies collected in Supplementary Movie 1 recorded with FER2-ckD parasites ±ATc in the presence of host cells. The TgFER2 replete parasite marked with the asterisk invades the host cell at the arrowhead. Invasion is complete in 20 seconds. The TgFER2-depleted parasite marked with asterisk makes an impulse move to the arrowhead and appears to deform the host cell. However, the parasite does not invade and disengages from the host cell reversing the deformation in the 40 sec frame. **(B)** Velocity profiles of FER2-ckD parasites ±ATc. The red arrowhead marks the synchronized frame where the parasites -ATc invade, or the parasites +ATc engage the host cell. Each thin line represents a single parasite from a single movie, heavy lines represent mean values for all parasites in each group included in the graph. Note that both sets of parasites show an impulse in motility right before the point of invasion/engagement, followed by an immediate pause, but that only the TgFER2 replete parasites maintain a positive velocity during the actual host cell invasion (magnified in the insert).

### TgFER2 is required for rhoptry secretion

As shown in Fig. 3, 7 and S7 the localization and morphology of the rhoptries were not affected by TgFER2 depletion. To test rhoptry function, we first monitored the release of rhoptry neck proteins by tracking RON4 distribution and assaying MJ formation (Fig. 7A). In the non-induced control we readily observed MJ formation whereas no MJ formation was detected in absence of TgFER2. These data suggest that the RON proteins are not secreted, or if they are, the do not assemble into the MJ.

**FIG 7.**
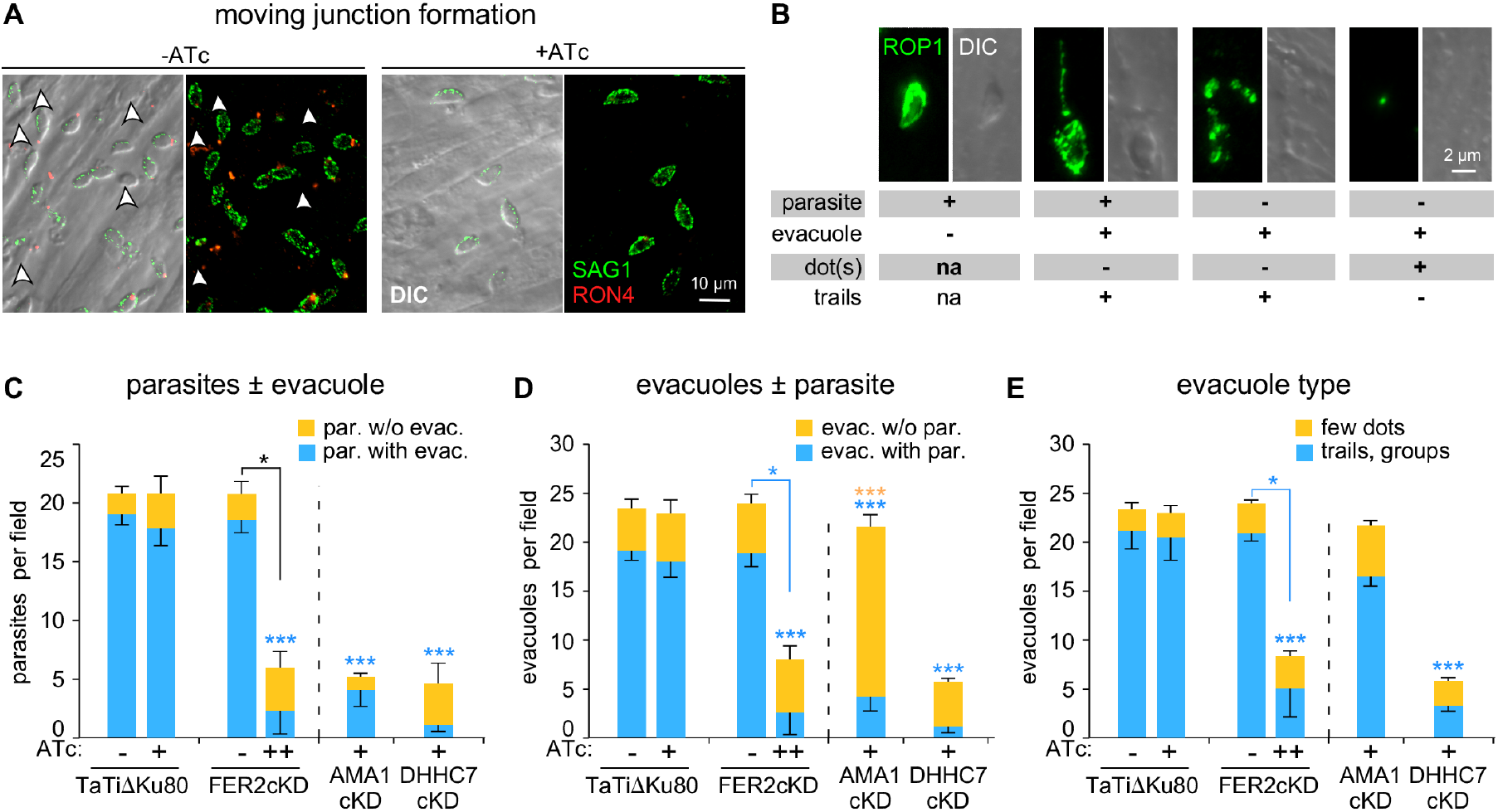
Rhoptry secretion of TgFER2 depleted parasites is impaired. **(A)** Formation of the MJ. Parasites were incubated with host cells for 10 min. MJ formation was visualized with RON4 antiserum under semi-permeabilizing conditions by 0.02% saponin. SAG1 stains the extracellular portion of the parasites. Arrowheads mark successfully invaded parasites that are not accessible to the SAG1 antibodies. Brightness and contrast adjustments are made identical for both conditions and thus signals are directly comparable. **(B)** Representative examples of parasite and evacuole features scored in the evacuole assay represented in panels C-E. na = not applicable. **(C-E)** Evacuole assay to monitor rhoptry bulb secretion and assess stability of the MJ attachment. Parasites as indicated were grown under ATc for 48 hr (+) or 96 hr (++) and incubated with host cells for 10 min. Evacuole formation was visualized using ROP1 antiserum following paraformaldehyde fixation. n=3, + or – stdev; Across samples statistics – or + ATc: one way ANOVA correction marked above bar. Pairwise ±ATc statistics marked above connector line; Student’s *t*-test. * P<0.01, ** P=0.001, *** P<0.0001. Asterisk color corresponds with the variable compared across samples; black asterisks correspond with analysis on the total number of parasites. The data presented in these panels are derived from the same experiments.

We next performed evacuole assays to determine the quantity and quality of rhoptry protein secretion in TgFER2 mutants. We used the AMA1-cKD as control for partial rhoptry secretion and weak MJ strength (31, 32) and DHHC7-cKD parasites as control for defective rhoptry secretion (5). Evacuoles, which are rhoptry protein clusters injected in the host cell cystosol, were visualized with ROP1 antiserum and classified as illustrated in Fig. 7B (32). To examine the strength of the overall parasite-host cell interaction the number of parasites per field was counted and differentiated by whether they were associated with an evacuole (Fig. 7C). The number of evacuoles per field and whether they are associated with parasites is a measure of the strength of the MJ (Fig. 7D). Levels of rhoptry secretion were differentiated by the relative size of evacuole patterns (Fig. 7E): small punctate ROP1 staining indicates less secretion than long trails or clusters. For FER2-ckD parasites, the number of parasites per field was consistent with the observations from the attachment and invasion assays: depletion of TgFER2 decreased parasite attachment (Fig. 7C). Among the parasites that were attached, very few were associated with evacuoles, indicating that they have not secreted the contents of their rhoptries into the host cell. The TgFER2 data are comparable with the results for the DHHC7 mutant. However, they differed from parasites that lack AMA1, where few parasites attach but the majority of the parasites have secreted rhoptries and generated evacuoles. TgFER2-depleted parasites, like the DHHC7 mutant, appear to secrete very few, if any, rhoptries. When we assess how many of the observed evacuoles are associated with parasites it becomes clear that TgFER2 depletion is much more similar to DHHC7 depletion than to parasites lacking AMA1 (Fig. 7D). Finally, we observed relatively few extensive evacuole patterns for both TgFER2 and DHHC7-depleted parasites (Fig. 7E) and conclude that in the rare events of rhoptry secretion, very little material was released. Overall, we conclude that TgFER2 is required for the secretion of the rhoptries, which is necessary to invade host cells.

## Discussion

Micronemes and rhoptries are essential to the invasion of apicomplexan parasites. These fascinating cellular structures are likely derived from ancestral organelles that persist in modern predatory protozoa and were adapted during the evolution of the Apicomplexa’s intracellular, parasitic lifestyle (9). While the molecular details of the initiation of microneme secretion are incompletely understood, the critical role of intracellular Ca^2+^ fluxes has been known for decades. More recently, intermediate players in the transduction of this signal (e.g. CDPKs and calcineurin) have been identified and the unconventional trafficking of microneme contents through a modified endosomal system has been described (36, 37). Far less is known about either the mechanisms of rhoptry secretion or the trafficking of their contents.

It has long been hypothesized that secretion from both micronemes and/or rhoptries requires a membrane fusion event, but evidence for a canonical secretion machinery has been elusive. Using C2 domains as the anchor for a bioinformatic search for potential components of this pathway, we were unable to find homologs for either synaptotagmins or the canonical DOC2 family of Ca^2+^ sensors function in mammalian neurotransmitter release (23, 38). We did find orthologs of the ferlin family of Ca^2+^ sensing membrane fusion proteins. TgFER1 and TgFER2 are widely conserved across the Apicomplexa, whereas the degenerate TgFER3 is found only in the Coccidia and a single Chromerid species, illustrative of the ancient history of these processes.

Detailed studies of *Toxoplasma* FER2 demonstrated it is required for secretion from the rhoptries. This finding provides one of the first mechanistic insights into rhoptry secretion, firmly linking it to the activity of this C2 domain-containing protein. It is generally accepted that rhoptry secretion must be preceded by microneme secretion and requires contact with an appropriate host cell (4, 8). However, neither the transduction of this attachment signal nor the process by which it leads to secretion of the organelle’s contents have been clarified. Although it is known that Mic8 is required for rhoptry release and has been postulated to be key in a signal transduction pathway (39), there is no experimental data supporting this model. Furthermore, AMA1 (32) and RON5 (40) also appear to be involved in rhoptry secretion but these mechanisms are equally unknown. Our finding that the Ca^2+^ sensor TgFER2 is required for rhoptry secretion provides a tantalizing hint at a more detailed mechanism. Although we have not been able to definitively demonstrate a role for Ca^2+^ in TgFER2 function, ferlins are Ca^2+^-sensing proteins and it is well established that a raise in [Ca^2+^]i accompanies host cell invasion (2). While the conventional belief has been that this fluctuation acted only on activation of motility, conoid extrusion and microneme secretion, we provide a hint that rhoptry secretion may be similarly dependent on variations in [Ca^2+^]_i_. The presence of an Asp residues constellation consistent with Ca^2+^-binding capacity in TgFER2’s C2F domain supports this model (see Fig S8 in the supplementary material), though the relative importance of the protein’s individual C2 domains in this process remain to be experimentally determined.

Of the mammalian ferlins, otoferlin is currently the best studied, yet its mechanism of action remains poorly understood (41). Otoferlin is expressed in many tissues, but in cochlear hair cells (CHCs) it controls the release of neurotransmitter upon an increase in [Ca^2+^]_i_ (42, 43). A raise in [Ca^2+^]_i_ leads otoferlin to interact with phospholipids (41) and SNARE proteins *in vitro* (44), though SNAREs have been debated to be absent from the site of secretion in CHCs (45). This highlights the potential for ferlin proteins to facilitate membrane fusion in the absence of SNAREs, an important parallel to *Toxoplasma*, in which there is currently no evidence for either rhoptry- or microneme-resident SNAREs. Otoferlin localizes to synaptic vesicles and the plasma membrane in CHCs (43). Our IEM observations on TgFER2 are consistent with a role in rhoptry secretion: in intracellular parasites TgFER2 localizes in a patchy pattern to the cytoplasmic side of the IMC next to a strong TgFER2 concentration inside the conoid; in extracellular parasites TgFER2 is detected in the conoid and surface of the rhoptries. This membrane transition is conceivable with Ca^2+^-dependent process (e.g. Ca^2+^-dependent phosphorylation (46)) and/or a change in membrane lipid composition of the IMC or rhoptry (e.g. as described for microneme secretion (47)).

As part of this study we compared different invasion and egress mutants across several commonly used assays, which allowed for several important observations. First, we observed that microneme protein mediated interactions are responsible for 50% of the attachment to fixed host cells. Somewhat unexpectedly, the red-green invasion assay did not differentiate the various mutants very well, with the exception of confirming the partial attachment defect previously demonstrated for the CnA mutant (11). Thus, this assay is not capable of specifically attributing individual phenotypes to defects in attachment versus invasion. By contrast, the evacuole assay was very powerful in differentiating different aspects of MJ formation and rhoptry secretion.

Overall, our findings support two interesting hypotheses. First, if ferlins act as Ca^2+^-sensors during Ca^2+^-dependent secretion in the Apicomplexa, TgFER2 may represent the link between the previously observed Ca^2+^ fluctuations during invasion and the well-described mechanics of MJ formation. If on the other hand the essential role of TgFER2 during rhoptry secretion is calcium-independent, this would signify a fascinating evolutionary divergence from the canonical function of ferlins as Ca^2+^ sensors. While additional work will be required to distinguish between these models, the work presented here is a critical step in our understanding of these critical virulence processes.

## Materials and Methods

### Parasites and mammalian cell lines

Transgenic derivatives of the RH strain the were maintained in human foreskin fibroblasts (HFF) as previously described (48). For the attachment assay under fluidics shear stress, HUVEC were cultured in EGM-2 medium containing EGM-2 SingleQuot supplements and growth factors (Lonza, Allendal, NJ). TgFER2 CDS was amplified using primers YFP-FER2-F/R and *NheI/EcoRV* cloned into tub-YFPYFP(MCS)/sagCAT (49) to generate ptub-YFP-FER2/sagCAT which was used for Sanger sequencing validation of the gene model. FER2-cKD parasites were generated by *BglīUNotI* cloning PCR amplified FER2 sequence (primers BamHI-FER2-F/NotI-FER2-R) into N-terminal myc-epitope tagged plasmid derived from DHFR-TetO7sag4-Nt-GOI (Wassim Daher, Université de Montpellier) and linearized by *XbaI* prior to transfection. ts-DOC2 parasites were generated by first 5xTY tagging the DOC2 locus using PCR amplicon (primers 5xTy_upstream_F/5xTy_PlusLink_R) from plasmid pLIC-5xTY-DD24/HX (Chris Tonkin, Walter and Eliza Hall Institute) and *Bgl*II/*Eco*RV cloning into tub-YFPYFP(MCS)/sagCAT. The tub promoter was *PmeI/BgľII* replaced with the 3’DOC2 homologous region PCR amplified from gDNA (primers DOC2_3-target_F/R). The CAT cassette was *PmeI/NotI* replaced with a DHFR minigene cassette and plasmid NheI linearized prior to transfection. A CRISPR/Cas9 plasmid was generated to mutate DOC2 F124 to S124 using primers DOC2_proto_F/R (50) and co-transfected with hybridized oligos DOC2_FM>SV_F/R in RHAKu80AHX-DOC2-5xTY parasites. All primer sequences are provided in Table S1 in the supplemental material.

### Imaging

The following antisera were used: α-Myc MAb 9E10, α-SAG1 MAb DG52 (51), α-Mic2 MAb 6D10 (52), mouse α-AMA1 (32), rabbit α-Mic3 (29) rabbit α-Mic5 (53), rabbit α-Mic8 (54), and mouse α-ROP1 (55). Alexa 488 or 594 conjugated secondary antibodies were used (Invitrogen). Images were collected on a Zeiss Axiovert 200 M wide-field fluorescence microscope and images were deconvolved and adjusted for phase contrast using Volocity software (Perkin Elmer).

### Egress assay

Assayed as described previously (11, 16). Freshly lysed, parasites, pretreated ±ATc for 24 hr were inoculated in HFF cells and incubated ±ATc for additional 24 hr. For 96 hr, parasites treated ±ATc for 68 hr were inoculated and incubated ±ATc for additional 30 hr. Egress was triggered by treatment with 2 μM A23187 or DMSO at 37°C for 5 min, followed by IFA with rat α-IMC3 (49). Intact vacuoles were counted for each sample in at least 10 fields and percentage egress calculated relative to the DMSO control

### Attachment and invasion

The combined attachment/invasion assay was performed as previously published (16, 33) with modifications described in (11): Tachyzoites treated ±ATc for the hrs as indicated (*ts*-DOC2 parasites incubated at 35°C and 40°C) were added to host cells in a 96-well plate, centrifuged (28*g, 3 min, RT), and allowed to invade for 1 hr at 37 C. Non-invaded extracellular parasites were detected using A594 conjugated α-SAG1 T41E5 (56). Following fixation and permeabilization, all parasites were visualized with A488 conjugated α-SAG1 T41E5. At least 300 parasites were counted per sample.

### Attachment to fixed host cells

Assay was performed as previously described (32). HFF confluent 96 well optical bottom plates were fixed with 3% formaldehyde + 0.06% glutaraldehyde for 5 min at 4°C, followed by overnight 0.16 M ethanolamine quenching at 4°C. Wells were pre-rinsed with 0.2% BSA in DMEM. Cytoplasmic YFP expressing TATiΔKu80 parasites mixed in 1:1 ratio were used as internal control (11), centrifuged (28*g, 5 min, 20°C) on the monolayer and incubated for 30 min at 37°C. Wells were rinsed 3 times with PBS, fixed with 4% PFA for 30 min at 4°C and permeabilized with 0. 25% TX-100 for 10 min. After blocking with 1% BSA in PBS, the parasites were probed with rabbit α-GFP (Torrey Pines Biolabs), and mouse α-SAG1 DG52. Three random fields in 3 independent wells were counted.

### Attachment under fluidics shear stress

was performed as described previously (34, 57). Microfluidic channels containing fibronectin were coated overnight with HUVEC. Freshly lysed parasites treated ±ATc for 48 or 96 hr were either stained with CMTPX CellTracker red or CFSE (Life Technologies), counted and combined 1:1. In each replicate experiment, the dyes were switched on the parental and knock-down parasite lines. Parasites were flowed at a shear force of 0.5 dyne/cm^2^ for 10 min at 37°C, and were fixed under flow conditions with 4% PFA for 30 min, followed by imaging on a Nikon Eclipse Ti microscope.

### Conoid extrusion assay

was performed as published (26). Freshly lysed parasites grown ±ATc for 48 hr were resuspended in 10% FBS in HS buffer. Conoid extrusion was induced using 0.5 M ethanol or 5 μM A23187 for 30 seconds. Parasites were fixed and scored for conoid extrusion by phase contrast microscopy. Samples were counted blindly scoring more than 350 parasites per sample.

### Microneme Mic2, Mic8, Mic10 secretion by western blot

was performed as published (30). Freshly lysed parasites treated ±ATc for 48 hr, resuspended in DMEM/FBS were added to a 96-well round-bottom plate and secretion induced by 1 μM A23187 or DMSO for 5 min at 37°C. Constitutive microneme secretion: no stimulation 37°C for 60 min. Supernatants were probed by western blot using MAb 6D10 α-Mic2 (52), rabbit α-Mic8 (54), rabbit α-Mic10 (58), and MAb α-Gra1 (59). Signals were quantified using a densitometer.

### Microneme Mic3, Mic5, Mic8 secretion by IFA

Mic3 (29), Mic5 (53), or Mic8 (54) IFA on parasites exposed to a host cell monolayer was performed as published (30). Parasites resuspended in Endo buffer were spun onto HFF cells in a 6-well plate (28*g, 5 min, RT) and incubated at 37°C for 20 min. Endo buffer was replaced by DMEM, 3% FBS and 10 mM HEPES pH7.2 and incubated at 37°C for 5 min. PBS washed coverslips were fixed with 4% formaldehyde / 0.02% glutaraldehyde followed by IFA in the presence of 0.02% saponin.

### Motility assessments

Motility was analyzed by video microscopy essentially as described previously (16). Intracellular tachyzoites grown for 96 hr ±ATc were physically harvested and resuspended in modified Ringer’s Medium and added to HFF confluent glass-bottom culture dishes (MatTek). The dish was imaged using a 63x objective at 37°C. Videos were recorded with 1 sec intervals. Velocities of individual invasion events were analyzed using the ImageJ/FIJI Cell Counter plug-in.

### Moving Junction (MJ) formation

was determined as published (39) with described modifications (11). Parasites grown ±ATc for 48 hr were inoculated into a HFF-confluent 24-well plate by centrifugation (28, 5 min, 20°C) and incubation at 37°C for 10 min. Wells were rinsed twice with PBS and fixed with 4% PFA at 4°C and partly permeabilized with 0.02% saponin. MJ was detected using rabbit α-RON4 (7) and all parasites were detected following full permeabilization with using MAb α-SAG1 DG52.

### Evacuole assay

Evacuoles were determined as described (60) with modifications. 1×10^7^ parasites grown ±ATc for 48 hr were inoculated into HFF confluent 24-well plates. The plate was centrifuged (28*g, 15 min, 23°C) and incubated at 37°C for 10 min. Wells were rinsed twice with PBS and fixed with 4% PFA at 4°C and 0.25% TX-100 permeabilized. Evacuoles were detected by MAb Tg49 α-ROP1 (55). More than 100 events per sample per experiment were counted.

### Immuno electron microscopy

Following washing with PBS, overnight infected HFF cells were fixed in 4% PFA in 0.25 M HEPES (pH 7.4) for 1 hr at RT, then in 8% PFA in the same buffer overnight at 4 C. They were infiltrated, frozen and sectioned as previously described (61). Sections were immunolabeled with α-Myc 9E10 in 1% fish skin gelatin, then with goat ±-IgG antibodies, followed by 10 nm protein A-gold particles before examination with a Philips CM120 electron microscope under 80 kV.

### Transmission electron microscopy

Parasites were fixed in 4% glutaraldehyde in 0.1 M phosphate buffer pH 7.4 and processed for routine electron microscopy (62). Briefly, cells were post-fixed in osmium tetroxide, and treated with uranyl acetate prior to dehydration in ethanol, treatment with propylene oxide, and embedding in Spurr’s epoxy resin. Thin sections were stained with uranyl acetate and lead citrate prior to examination with a JEOL 1200EX electron microscope.

### Statistics

Student’s paired *t*-test and one-way ANOVA using posthoc Bonferroni correction were used where indicated against the TaTiΔKu80 line.

## Acknowledgements

We thank Dr. Jamie Henzy for assistance with phylogenetic analysis, Dr. Sander Groffen for assistance with modeling, Amir Bayegan for assistance with statistics, Elizabeth C. Gray for technical assistance, Drs. Bradley, Boothroyd, Carruthers, Cesbron-Delauw, Daher, de Graffenried, Dubremetz, Saeij, Sibley, Soldati, Striepen, Tonkin, and Ward for sharing reagents, and Dr Manoj Duraisingh for critically reading the manuscript. We would also like to thank the technical competence of Kimberley Zichichi from the Electron Microscopy Facility at Yale University.

This study was supported by NIH AI108251 (BIC), AI060767 (IC), AI099658 (MJG), and AI122923 (MJG), Deutsche Forschungsgemeinschaft (KE), American Cancer Society 126688-RSG-14-202-01-MPC (MBL) and American Cancer Society RSG-12-175-01-MPC (MJG). The funders had no role in study design, data collection and analysis, decision to publish, or preparation of the manuscript.

## References

1. Kafsack BF, Pena JD, Coppens I, Ravindran S, Boothroyd JC, Carruthers VB. 2009. Rapid membrane disruption by a perforin-like protein facilitates parasite exit from host cells. Science 323:530–3.

2. Wetzel DM, Chen LA, Ruiz FA, Moreno SN, Sibley LD. 2004. Calcium-mediated protein secretion potentiates motility in Toxoplasma gondii. J Cell Sci 117:5739–48.

3. Carruthers VB, Tomley FM. 2008. Microneme proteins in apicomplexans. Subcell Biochem 47:33–45.

4. Carruthers V, Boothroyd JC. 2007. Pulling together: an integrated model of Toxoplasma cell invasion. Curr Opin Microbiol 10:83–9.

5. Beck JR, Fung C, Straub KW, Coppens I, Vashisht AA, Wohlschlegel JA, Bradley PJ. 2013. A Toxoplasma palmitoyl acyl transferase and the palmitoylated Armadillo Repeat protein TgARO govern apical rhoptry tethering and reveal a critical role for the rhoptries in host cell invasion but not egress. PLoS Pathog 9:e1003162.

6. Bradley PJ, Sibley LD. 2007. Rhoptries: an arsenal of secreted virulence factors. Curr Opin Microbiol 10:582–7.

7. Alexander DL, Mital J, Ward GE, Bradley P, Boothroyd JC. 2005. Identification of the Moving Junction Complex of Toxoplasma gondii: A Collaboration between Distinct Secretory Organelles. PLoS Pathog 1:e17.

8. Carruthers VB, Sibley LD. 1997. Sequential protein secretion from three distinct organelles of Toxoplasma gondii accompanies invasion of human fibroblasts. Eur J Cell Biol 73:114–23.

9. Gubbels MJ, Duraisingh MT. 2012. Evolution of apicomplexan secretory organelles. Int J Parasitol 42:1071–81.

10. Mercier C, Cesbron-Delauw MF. 2015. Toxoplasma secretory granules: one population or more? Trends Parasitol doi:10.1016/j.pt.2014.12.002.

11. Paul AS, Saha S, Engelberg K, Jiang RH, Coleman BI, Kosber AL, Chen CT, Ganter M, Espy N, Gilberger TW, Gubbels MJ, Duraisingh MT. 2015. Parasite Calcineurin Regulates Host Cell Recognition and Attachment by Apicomplexans. Cell Host Microbe 18:49–60.

12. Lourido S, Shuman J, Zhang C, Shokat KM, Hui R, Sibley LD. 2010. Calcium-dependent protein kinase 1 is an essential regulator of exocytosis in Toxoplasma. Nature 465:359–62.

13. Garrison E, Treeck M, Ehret E, Butz H, Garbuz T, Oswald BP, Settles M, Boothroyd J, Arrizabalaga G. 2012. A forward genetic screen reveals that calcium-dependent protein kinase 3 regulates egress in Toxoplasma. PLoS Pathog 8:e1003049.

14. Lourido S, Tang K, Sibley LD. 2012. Distinct signalling pathways control Toxoplasma egress and host-cell invasion. Embo J 31:4524–34.

15. McCoy JM, Whitehead L, van Dooren GG, Tonkin CJ. 2012. TgCDPK3 Regulates Calcium-Dependent Egress of Toxoplasma gondii from Host Cells. PLoS Pathog 8:e1003066.

16. Farrell A, Thirugnanam S, Lorestani A, Dvorin JD, Eidell KP, Ferguson DJ, Anderson-White BR, Duraisingh MT, Marth GT, Gubbels MJ. 2012. A DOC2 protein identified by mutational profiling is essential for apicomplexan parasite exocytosis. Science 335:218–21.

17. Martens S, McMahon HT. 2008. Mechanisms of membrane fusion: disparate players and common principles. Nat Rev Mol Cell Biol 9:543–56.

18. Martens S. 2010. Role of C2 domain proteins during synaptic vesicle exocytosis. Biochem Soc Trans 38:213–6.

19. Lek A, Lek M, North KN, Cooper ST. 2010. Phylogenetic analysis of ferlin genes reveals ancient eukaryotic origins. BMC Evol Biol 10:231.

20. Lek A, Evesson FJ, Sutton RB, North KN, Cooper ST. 2012. Ferlins: regulators of vesicle fusion for auditory neurotransmission, receptor trafficking and membrane repair. Traffic 13:185–94.

21. Pang ZP, Sudhof TC. 2010. Cell biology of Ca2+-triggered exocytosis. Curr Opin Cell Biol 22:496–505.

22. Sudhof TC. 2013. A molecular machine for neurotransmitter release: synaptotagmin and beyond. Nat Med 19:1227–31.

23. Walter AM, Groffen AJ, Sorensen JB, Verhage M. 2011. Multiple Ca2+ sensors in secretion: teammates, competitors or autocrats? Trends Neurosci 34:487–97.

24. Woo YH, Ansari H, Otto TD, Klinger CM, Kolisko M, Michalek J, Saxena A, Shanmugam D, Tayyrov A, Veluchamy A, Ali S, Bernal A, Del Campo J, Cihlar J, Flegontov P, Gornik SG, Hajduskova E, Horak A, Janouskovec J, Katris NJ, Mast FD, Miranda-Saavedra D, Mourier T, Naeem R, Nair M, Panigrahi AK, Rawlings ND, Padron-Regalado E, Ramaprasad A, Samad N, Tomcala A, Wilkes J, Neafsey DE, Doerig C, Bowler C, Keeling PJ, Roos DS, Dacks JB, Templeton TJ, Waller RF, Lukes J, Obornik M, Pain A. 2015. Chromerid genomes reveal the evolutionary path from photosynthetic algae to obligate intracellular parasites. Elife 4.

25. Meissner M, Schluter D, Soldati D. 2002. Role of Toxoplasma gondii myosin A in powering parasite gliding and host cell invasion. Science 298:837–40.

26. Mondragon R, Frixione E. 1996. Ca(2+)-dependence of conoid extrusion in Toxoplasma gondii tachyzoites. J Eukaryot Microbiol 43:120–7.

27. Carruthers VB, Moreno SN, Sibley LD. 1999. Ethanol and acetaldehyde elevate intracellular [Ca2+] and stimulate microneme discharge in Toxoplasma gondii. Biochem J 342 (Pt 2):379–86.

28. Kremer K, Kamin D, Rittweger E, Wilkes J, Flammer H, Mahler S, Heng J, Tonkin CJ, Langsley G, Hell SW, Carruthers VB, Ferguson DJ, Meissner M. 2013. An overexpression screen of Toxoplasma gondii Rab-GTPases reveals distinct transport routes to the micronemes. PLoS Pathog 9:e1003213.

29. Garcia-Reguet N, Lebrun M, Fourmaux MN, Mercereau-Puijalon O, Mann T, Beckers CJ, Samyn B, Van Beeumen J, Bout D, Dubremetz JF. 2000. The microneme protein MIC3 of Toxoplasma gondii is a secretory adhesin that binds to both the surface of the host cells and the surface of the parasite. Cell Microbiol 2:353–64.

30. Carruthers VB, Sibley LD. 1999. Mobilization of intracellular calcium stimulates microneme discharge in Toxoplasma gondii. Mol Microbiol 31:421–8.

31. Lamarque MH, Roques M, Kong-Hap M, Tonkin ML, Rugarabamu G, Marq JB, Penarete-Vargas DM, Boulanger MJ, Soldati-Favre D, Lebrun M. 2014. Plasticity and redundancy among AMA-RON pairs ensure host cell entry of Toxoplasma parasites. Nat Commun 5:4098.

32. Mital J, Meissner M, Soldati D, Ward GE. 2005. Conditional Expression of Toxoplasma gondii Apical Membrane Antigen-1 (TgAMA1) Demonstrates That TgAMA1 Plays a Critical Role in Host Cell Invasion. Mol Biol Cell 16:4341–9.

33. Carey KL, Westwood NJ, Mitchison TJ, Ward GE. 2004. A small-molecule approach to studying invasive mechanisms of Toxoplasma gondii. Proc Natl Acad Sci U S A 101:7433–8.

34. Harker KS, Jivan E, McWhorter FY, Liu WF, Lodoen MB. 2014. Shear forces enhance Toxoplasma gondii tachyzoite motility on vascular endothelium. mBio 5:e01111–13.

35. Bichet M, Joly C, Hadj Henni A, Guilbert T, Xemard M, Tafani V, Lagal V, Charras G, Tardieux I. 2014. The toxoplasma-host cell junction is anchored to the cell cortex to sustain parasite invasive force. BMC Biol 12:773.

36. Sangare LO, Alayi TD, Westermann B, Hovasse A, Sindikubwabo F, Callebaut I, Werkmeister E, Lafont F, Slomianny C, Hakimi MA, Van Dorsselaer A, Schaeffer-Reiss C, Tomavo S. 2016. Unconventional endosome-like compartment and retromer complex in Toxoplasma gondii govern parasite integrity and host infection. Nat Commun 7:11191.

37. Tomavo S. 2014. Evolutionary repurposing of endosomal systems for apical organelle biogenesis in Toxoplasma gondii. Int J Parasitol 44:133–8.

38. Shin OH. 2014. Exocytosis and synaptic vesicle fusion. Comprehensive Physiology 4:149–175.

39. Kessler H, Herm-Gotz A, Hegge S, Rauch M, Soldati-Favre D, Frischknecht F, Meissner M. 2008. Microneme protein 8–a new essential invasion factor in Toxoplasma gondii. J Cell Sci 121:947–56.

40. Beck JR, Chen AL, Kim EW, Bradley PJ. 2014. RON5 is critical for organization and function of the Toxoplasma moving junction complex. PLoS Pathog 10:e1004025.

41. Pangrsic T, Reisinger E, Moser T. 2012. Otoferlin: a multi-C2 domain protein essential for hearing. Trends in neurosciences 35:671–80.

42. Roux I, Safieddine S, Nouvian R, Grati M, Simmler MC, Bahloul A, Perfettini I, Le Gall M, Rostaing P, Hamard G, Triller A, Avan P, Moser T, Petit C. 2006. Otoferlin, defective in a human deafness form, is essential for exocytosis at the auditory ribbon synapse. Cell 127:277–89.

43. Pangrsic T, Lasarow L, Reuter K, Takago H, Schwander M, Riedel D, Frank T, Tarantino LM, Bailey JS, Strenzke N, Brose N, Muller U, Reisinger E, Moser T. 2010. Hearing requires otoferlin-dependent efficient replenishment of synaptic vesicles in hair cells. Nat Neurosci 13:869–76.

44. Johnson CP, Chapman ER. 2010. Otoferlin is a calcium sensor that directly regulates SNARE-mediated membrane fusion. J Cell Biol 191:187–97.

45. Nouvian R, Neef J, Bulankina AV, Reisinger E, Pangrsic T, Frank T, Sikorra S, Brose N, Binz T, Moser T. 2011. Exocytosis at the hair cell ribbon synapse apparently operates without neuronal SNARE proteins. Nat Neurosci 14:411–3.

46. Meese S, Cepeda AP, Gahlen F, Adams CM, Ficner R, Ricci AJ, Heller S, Reisinger E, Herget M. 2017. Activity-Dependent Phosphorylation by CaMKIIdelta Alters the Ca(2+) Affinity of the Multi-C2-Domain Protein Otoferlin. Front Synaptic Neurosci 9:13.

47. Bullen HE, Jia Y, Yamaryo-Botte Y, Bisio H, Zhang O, Jemelin NK, Marq JB, Carruthers V, Botte CY, Soldati-Favre D. 2016. Phosphatidic Acid-Mediated Signaling Regulates Microneme Secretion in Toxoplasma. Cell Host Microbe 19:349–60.

48. Roos DS, Donald RG, Morrissette NS, Moulton AL. 1994. Molecular tools for genetic dissection of the protozoan parasite Toxoplasma gondii. Methods Cell Biol 45:27–63.

49. Anderson-White BR, Ivey FD, Cheng K, Szatanek T, Lorestani A, Beckers CJ, Ferguson DJ, Sahoo N, Gubbels MJ. 2011. A family of intermediate filament-like proteins is sequentially assembled into the cytoskeleton of Toxoplasma gondii. Cell Microbiol 13:18–31.

50. Sidik SM, Hackett CG, Tran F, Westwood NJ, Lourido S. 2014. Efficient genome engineering of Toxoplasma gondii using CRISPR/Cas9. PLoS ONE 9:e100450.

51. Burg JL, Perelman D, Kasper LH, Ware PL, Boothroyd JC. 1988. Molecular analysis of the gene encoding the major surface antigen of Toxoplasma gondii. Journal of immunology 141:3584–91.

52. Wan KL, Carruthers VB, Sibley LD, Ajioka JW. 1997. Molecular characterisation of an expressed sequence tag locus of Toxoplasma gondii encoding the micronemal protein MIC2. Mol Biochem Parasitol 84:203–14.

53. Brydges SD, Zhou XW, Huynh MH, Harper JM, Mital J, Adjogble KD, Daubener W, Ward GE, Carruthers VB. 2006. Targeted deletion of MIC5 enhances trimming proteolysis of Toxoplasma invasion proteins. Eukaryot Cell 5:2174–83.

54. Meissner M, Reiss M, Viebig N, Carruthers VB, Toursel C, Tomavo S, Ajioka JW, Soldati D. 2002. A family of transmembrane microneme proteins of Toxoplasma gondii contain EGF-like domains and function as escorters. J Cell Sci 115:563–74.

55. Saffer LD, Mercereau-Puijalon O, Dubremetz JF, Schwartzman JD. 1992. Localization of a Toxoplasma gondii rhoptry protein by immunoelectron microscopy during and after host cell penetration. J Protozool 39:526–30.

56. Couvreur G, Sadak A, Fortier B, Dubremetz JF. 1988. Surface antigens of Toxoplasma gondii. Parasitology 97 (Pt 1):1–10.

57. Ueno N, Harker KS, Clarke EV, McWhorter FY, Liu WF, Tenner AJ, Lodoen MB. 2014. Real-time imaging of Toxoplasma-infected human monocytes under fluidic shear stress reveals rapid translocation of intracellular parasites across endothelial barriers. Cell Microbiol 16:580–95.

58. Hoff EF, Cook SH, Sherman GD, Harper JM, Ferguson DJ, Dubremetz JF, Carruthers VB. 2001. Toxoplasma gondii: molecular cloning and characterization of a novel 18-kDa secretory antigen, TgMIC10. Exp Parasitol 97:77–88.

59. Cesbron-Delauw MF, Guy B, Torpier G, Pierce RJ, Lenzen G, Cesbron JY, Charif H, Lepage P, Darcy F, Lecocq JP. 1989. Molecular characterization of a 23-kilodalton major antigen secreted by Toxoplasma gondii. Proc Natl Acad Sci U S A 86:7537–41.

60. Hakansson S, Charron AJ, Sibley LD. 2001. Toxoplasma evacuoles: a two-step process of secretion and fusion forms the parasitophorous vacuole. Embo J 20:3132–44.

61. Folsch H, Pypaert M, Schu P, Mellman I. 2001. Distribution and function of AP-1 clathrin adaptor complexes in polarized epithelial cells. J Cell Biol 152:595–606.

62. Ferguson DJ, Cesbron-Delauw MF, Dubremetz JF, Sibley LD, Joiner KA, Wright S. 1999. The expression and distribution of dense granule proteins in the enteric (Coccidian) forms of Toxoplasma gondii in the small intestine of the cat. Exp Parasitol 91:203–11.

63. Kearse M, Moir R, Wilson A, Stones-Havas S, Cheung M, Sturrock S, Buxton S, Cooper A, Markowitz S, Duran C, Thierer T, Ashton B, Meintjes P, Drummond A. 2012. Geneious Basic: an integrated and extendable desktop software platform for the organization and analysis of sequence data. Bioinformatics 28:1647–9.

